# CentroFinder: accurate *de novo* identification of centromeres in fungal genomes

**DOI:** 10.64898/2026.02.04.702907

**Authors:** Sahar Salimi, Sharon Colson, Michael Renfro, Li-Jun Ma, Mostafa Rahnama

## Abstract

**Motivation:** Centromeres are essential chromosomal loci, yet their computational identification remains challenging due to rapid sequence evolution, high repeat content, and the absence of conserved defining motifs. This challenge is particularly pronounced in fungi, where centromere architectures vary widely in size, sequence composition, and chromatin organization, limiting the effectiveness of single-feature or motif-based prediction approaches.

**Results:** We present CentroFinder, a fungal-specific computational framework for *de novo* centromere prediction from long-read sequencing–based genome assemblies. CentroFinder integrates multiple genomic and long-read–derived features into a weighted scoring model to identify loci where centromere-associated signals converge. Benchmarking against experimentally mapped centromeres in *Cryptococcus deuterogattii, Magnaporthe oryzae*, and *Neurospora crassa* demonstrates that CentroFinder consistently predicts a single centromeric region per chromosome, fully nested within CENP-A–defined domains despite substantial diversity in centromere size, sequence composition, and chromatin context.

**Availability and Implementation:** CentroFinder is freely available as open-source software at https://github.com/RahnamaLab/CentroFinder. The pipeline is designed for high-performance computing environments and leverages features derived from long-read sequencing data.

## 1 Introduction

Centromeres are indispensable chromosomal loci that orchestrate kinetochore assembly and enable the attachment of spindle microtubules, ensuring the equal segregation of sister chromatids during both mitosis and meiosis [1, 2]. Their essential role in maintaining genome stability is underscored by the fact that centromere malfunction is associated with aneuploidy, cancer, and developmental disorders [3-5]. Despite this deeply conserved function, the underlying DNA sequences of centromeres exhibit extraordinary diversity and rapid evolution—an observation described as the “centromere paradox” [6-8]. As a result, centromere identity is governed not by a universal sequence motif but by the interplay among DNA sequence, repetitive content, chromatin organization, and epigenetic regulation [1].

Centromeric and pericentromeric regions have long been among the most poorly resolved parts of eukaryotic genomes because their high repeat content hinders assembly and read mapping with short-read data [9, 10]. The advent of long-read sequencing technologies such as PacBio HiFi and Oxford Nanopore, along with improved assembly algorithms, now enables complete reconstruction of these complex loci in many organisms [11-14], opening the door to systematic, sequence-based centromere analysis. The fungal kingdom exemplifies extreme centromere diversity, ranging from ≤400 bp genetically defined point centromeres in *Saccharomyces cerevisiae* to large regional centromeres exceeding 150 kb [8, 15-17]. Because no single genomic feature universally defines centromeres across fungi—and even closely related species may vary drastically [17]—computational prediction remains challenging, particularly for large regional centromeres embedded in repetitive DNA.

Although several software tools exist for centromere prediction, none are specifically designed for fungi, and most rely on a single dominant feature such as DNA methylation or tandem repeats [18-20]. While such approaches may perform well in species with relatively uniform centromere architecture, they often achieve low recall in fungi because centromeres are highly heterogeneous and no single feature is universally present [15, 16, 18, 19]. This gap highlights the need for a multi-feature, fungal-tailored prediction framework. To address this challenge, we developed CentroFinder, a fungal-specific computational pipeline that integrates multiple genomic and epigenetic signals to identify centromere locations in chromosome-level assemblies. CentroFinder quantifies six key features—TE enrichment, tandem-repeat density, DNA methylation, gene absence, GC depletion, and long-read depth anomalies—across the genome and combines them using a biologically informed weighted scoring model. By integrating these complementary signals and aggregating high-scoring regions into continuous intervals, CentroFinder produces robust and accurate *de novo* centromere predictions, enabling comparative analyses of karyotype evolution, centromere repositioning, and structural variation across fungal genomes.

## 2 Materials and methods

### 2.1 Genome Assemblies and Input Data

All centromere predictions were performed using high-quality fungal genome assemblies generated from long-read sequencing technologies (PacBio HiFi or Oxford Nanopore). Because centromere identification depends strongly on assembly continuity, analyses were restricted to assemblies with long contiguous internal regions spanning centromeric loci, regardless of whether they were fully chromosome-level or partially contiguous. For genomes lacking published annotations, gene models were generated *de novo* using FunGAP [21] to ensure standardized annotation quality across species.

### 2.2 Feature extraction for centromere prediction

To identify centromeres, we quantified a set of sequence- and long-read–derived genomic features known to characterize fungal regional centromeres, including transposable element (TE) density, tandem repeat (TR) density, DNA methylation signatures, GC content, gene density, and long-read sequencing coverage. All features were calculated in non-overlapping 1-kb windows across each chromosome and used as inputs for the integrated centromere-scoring model.

#### 2.2.1 Transposable element annotation

TEs were annotated using Extensive de-novo TE Annotator (EDTA) with default parameters [22]. Coding sequences were removed prior to TE annotation to avoid misclassification of protein-coding regions. The resulting GFF3 annotations were converted to BED format and summarized to compute TE density along each chromosome.

#### 2.2.2 Tandem repeat detection

TRs were identified using Tandem Repeats Finder (TRF) [23]. TRF outputs were converted to BED format using trf2bed, and TR coverage was aggregated to generate tandem-repeat density profiles across chromosomes.

#### 2.2.3 DNA methylation detection from long-read sequencing

For genomes sequenced with PacBio HiFi, CpG methylation was computed using the ccsmeth workflow [24]. First, PacBio subreads were converted into CCS (HiFi) reads using *pbccs* as implemented in ccsmeth. CCS reads were then aligned to the genome using *pbmm2* (https://github.com/PacificBiosciences/pbmm2). CpG methylation (5mC) was detected using ccsmeth call_mods with the *attbigru2s_b21*.*v3* model, and window-level methylation frequencies were computed with ccsmeth call_freqb in aggregate mode. Resulting methylation-frequency tracks were exported in BED format for downstream integration.

For genomes sequenced with Oxford Nanopore, methylation was extracted from raw signal–derived base-modification tags embedded in aligned reads. Nanopore reads were aligned to each genome using a minimap2-based workflow, and the resulting BAM files were sorted and indexed [25]. Base-resolution 5mC calls were obtained by converting the BAM files into BED format using modbam2bed (https://github.com/epi2me-labs/modbam2bed), which parses modified-base tags from individual reads. The resulting BED tracks provided per-site methylation information that was subsequently summarized into the analysis windows used for feature extraction.

Although PacBio HiFi and Oxford Nanopore platforms rely on distinct sequencing chemistries and methylation-calling algorithms, both approaches yield quantitative CpG methylation signals that are comparable at the genomic window scale used in this study. Because CentroFinder integrates methylation as a relative feature—capturing local enrichment or depletion patterns rather than absolute methylation levels—platform-specific differences do not affect centromere localization. In all cases, methylation tracks were summarized into uniform 1-kb windows prior to integration, ensuring consistency across sequencing technologies.

#### 2.2.4 Long-read coverage profiling

HiFi or Nanopore reads were aligned to each corresponding assembly using *pbmm2* (PacBio) or *minimap2* (Nanopore) [25]. Per-base read depth was aggregated across the genome, and a depth-anomaly metric—defined as deviation from the chromosome-wide median coverage—was calculated to highlight regions enriched in structurally complex or low-mappability sequences.

#### 2.2.5 Gene-content and GC-content extraction

Gene coordinates from FunGap or published annotations were converted to BED format and intersected with genomic windows to compute gene counts. GC content for each window was calculated directly from the genome assembly using *bedtools nuc* [26]. Low gene density (including windows with zero genes) and reduced GC content were interpreted as signatures of gene absence and GC depletion, respectively. These features were included because fungal centromeres typically exhibit low GC content and are depleted of protein-coding genes.

### 2.3 Centromere scoring and prediction

Each chromosome was partitioned into non-overlapping 1-kb windows, and six features were computed for each window: TE coverage, tandem-repeat density, CpG methylation frequency, long-read coverage anomaly, gene density, and GC content. To ensure comparability across disparate data types, all feature values were normalized to a 0–1 scale prior to integration.

For each window, the Centromere Score was calculated using a weighted linear combination of these normalized features:

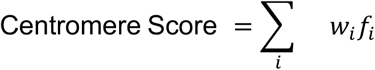

where *f*_*i*_ represents the normalized value of the *i*-th genomic feature for a given 1-kb window. Specifically, *f*_*i*_ corresponds to TE coverage, tandem-repeat density, CpG methylation frequency, long-read coverage anomaly, gene density, and GC content. Normalization of all *f*_*i*_ values to a 0–1 scale ensures that features measured in different units contribute comparably to the composite score and that feature weights, rather than raw signal magnitude, determine relative importance.

Feature weights (*w*) were assigned to reflect the known biological hallmarks of fungal regional centromeres. TRF density was prioritized with a weight of 4, and TE content with a weight of 3, as these represent the primary structural components of centromeric heterochromatin. The remaining features—reduced CpG methylation, low gene density (gene absence), reduced GC content (GC depletion), and long-read coverage anomaly— were each assigned a weight of 1. These parameters were selected to capture the characteristic “gene desert” and “repeat island” signature typical of fungal centromeres, where Repeat-Induced Point (RIP) mutations often lead to significant GC depletion. The weighting system is inherently flexible, allowing for adjustments based on the specific centromere architecture of the target species; for instance, weights may be shifted to prioritize methylation in species where tandem repeats are less pronounced.

Following score calculation, adjacent high-scoring windows were merged into continuous candidate intervals and the highest-scoring internal interval remaining after subtelomeric masking was designated as the predicted centromere for each chromosome. To prevent false-positive predictions in subtelomeric regions—typically repeat-rich, gene-poor, and structurally dynamic—windows within 500 kb of chromosome ends (for core chromosomes) or 50 kb (for accessory/mini-chromosomes) were excluded.

### 2.4 Benchmarking using genomes with experimentally validated centromeres

Pipeline performance was evaluated using a curated benchmarking dataset consisting of three fungal species with experimentally mapped centromeres (*Cryptococcus deuterogattii* R265 [27], *Magnaporthe oryzae* Guy11 [28], and *Neurospora crassa* FGSC2489 OR74A) [29]. For each species, predicted centromeres were compared against published centromere coordinates, and accuracy metrics were computed per chromosome and per species.

## 3 Results

### 3.1 CentroFinder Accurately Identifies Compact Regional Centromeres Across Fungal Genomes

CentroFinder reliably detected centromeres across chromosomes from three benchmark fungal species, each representing lineages with well-characterized regional centromeres [27-29]. Despite spanning diverse genome architectures, chromosome sizes, and repeat landscapes, the tool consistently identified a single candidate centromeric region per chromosome (Table S1). Across all genomes, CentroFinder recovered regions exhibiting the full suite of regional-centromere signatures, including GC depletion, enrichment of transposable elements and tandem repeats, reduced CpG methylation and gene density, and characteristic long-read coverage patterns. These hallmark properties collectively define regional centromeres in many fungi and provided a robust, independent validation of CentroFinder’s prediction strategy (Figure 1).

**Figure 1.**
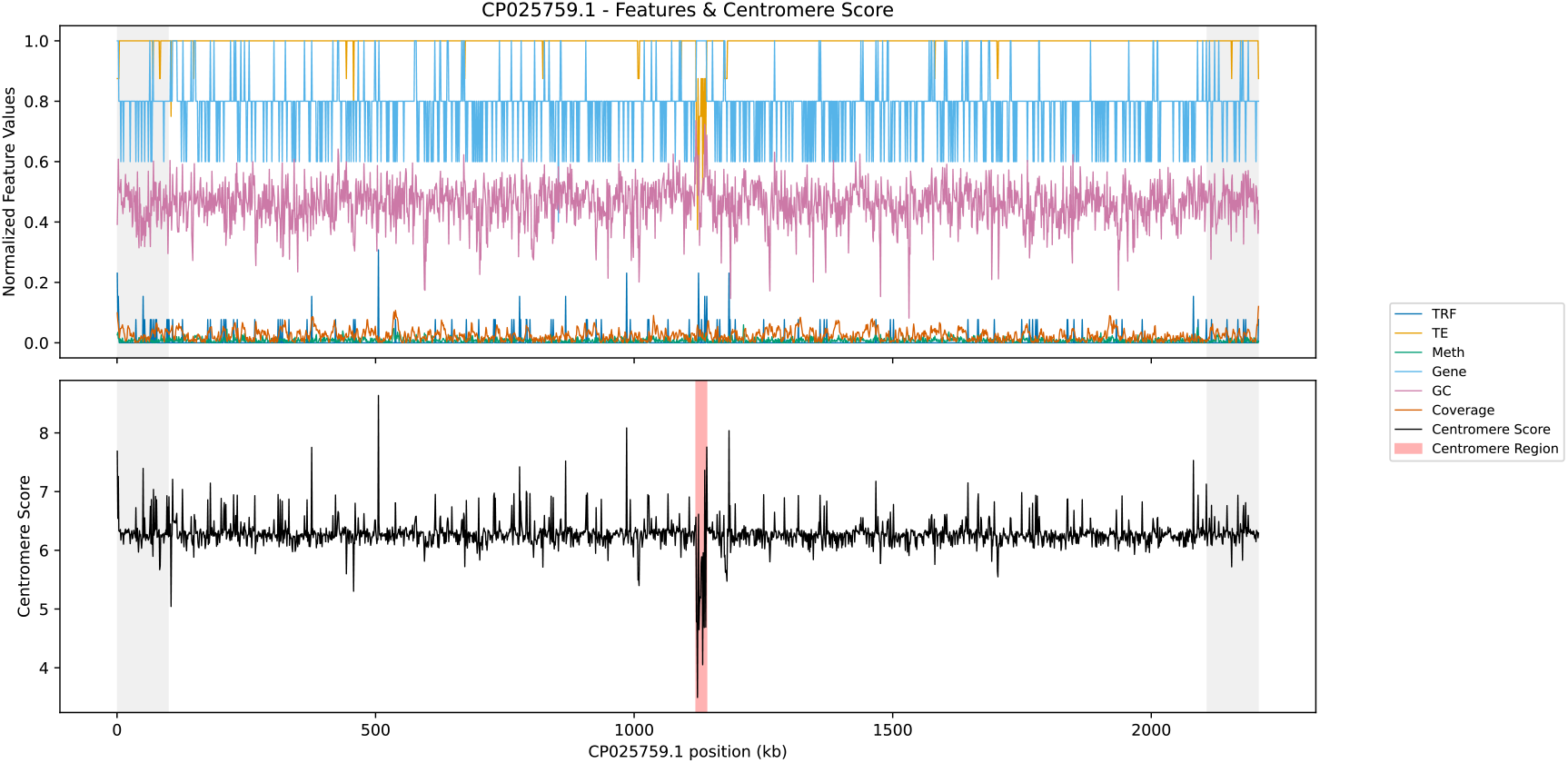
Representative CentroFinder output for chromosome CP025759.1 of *Cryptococcus deuterogattii*. The upper panel displays the linear profiles of the individual sequence- and long-read–derived features used by CentroFinder along the chromosome, including tandem repeat density (TRF), transposable element enrichment (TE), CpG methylation frequency (Meth), gene density (Gene), GC content (GC), and long-read coverage (Coverage). The lower panel shows the integrated CentroFinder score (black line), calculated as a weighted combination of these features across genomic windows. Gray-shaded regions at both chromosome ends indicate subtelomeric regions excluded from centromere prediction. The predicted centromeric region is highlighted by a light red box.

Across species, predicted centromere sizes varied in accordance with known centromere architectures but were consistently compact relative to experimentally defined centromeric domains. In *C. deuterogattii*, predicted centromeres typically spanned 4–20 kb, whereas larger predicted intervals were observed in *M. oryzae* (∼21–107 kb) and *N. crassa* (∼40–95 kb), reflecting species-specific centromere organization (Table S1). This pattern underscores the method’s ability not only to detect centromeres but also to consistently localize conserved centromeric cores within broader centromeric regions. Importantly, predictions were obtained using sequence- and long-read–derived features alone, without requiring chromatin immunoprecipitation or CENP-A mapping, demonstrating the tool’s utility for newly sequenced genomes or species where experimental mapping is not feasible.

### 3.2 High Concordance Between CentroFinder Predictions and Experimentally Defined Centromeres

To assess prediction accuracy, CentroFinder predictions were directly compared with experimentally defined centromeres (EDCs) derived from CENP-A or CenH3 mapping in three fungal species: *C. deuterogattii* R265, *M. oryzae* Guy11, and *N. crassa* OR74A. Across all evaluated chromosomes, CentroFinder identified exactly one centromeric region per chromosome that overlapped the corresponding experimentally defined centromere, demonstrating high positional accuracy and sensitivity (Table 1; Table S1). In *C. deuterogattii*, CentroFinder successfully predicted centromeres on all 14 chromosomes, with predicted intervals fully contained within the experimentally mapped centromeric regions (Table 1; Table S1). Positional offsets between predicted and experimental centromere boundaries ranged from 310 bp to 6.7 kb, with a median offset of approximately 1.1 kb (Table 1). Predicted centromere lengths were comparable to experimentally defined centromeric regions, typically spanning 4–20 kb compared with 8–22 kb for CENP-A–mapped domains, and were consistently nested within experimental boundaries.

**Table 1.** Concordance between CentroFinder predictions and experimentally defined centromeres in three fungal genomes. CentroFinder predictions were compared with experimentally defined centromeres (EDCs) derived from CENP-A or CenH3 mapping. Offsets represent the distance by which experimentally defined centromeric regions extend beyond the CentroFinder-predicted centromeric interval, calculated as the difference between corresponding start and end coordinates of the EDC and the prediction. Because predicted centromeres are typically fully contained within EDCs, offsets primarily reflect differences in boundary definitions rather than positional discordance. Median offset represents the median of offsets within each species.

| Species | No. of chromosomes evaluated | CentroFinder prediction length (kb) | EDC length (kb) | Offset range (kb) | Median offset (kb) |
| --- | --- | --- | --- | --- | --- |
| *Cryptococcus deuterogattii* | 14 | 4–20 | 8–22 | 0.31–6.7 | 1.1 |
| *Magnaporthe oryzae* | 7 | 21–107 | 57–109 | 0.16–36 | 26 |
| *Neurospora crassa* | 7<sup>†</sup> | 40–95 | 316–423 | 253–341 | 277 |

† Based on chromosomes with published EDCs

A similar pattern was observed in *M. oryzae*, where CentroFinder predictions overlapped experimentally mapped CenH3-associated regions on all chromosomes (Table 1; Table S1). Although experimentally defined centromeric regions in *M. oryzae* span substantially larger genomic intervals (∼57–109 kb), predicted centromeric regions were consistently smaller (∼21–107 kb) and largely nested within these domains. Positional offsets ranged from ∼0.16 kb to ∼36 kb, with a median offset of ∼26 kb (Table 1). On chromosome 5, the CentroFinder-predicted interval partially extended beyond the 3′ boundary of the experimentally defined centromere, while still overlapping the CenH3-enriched region. Overall, CentroFinder consistently delineates the compact, functionally relevant centromeric core, while broader experimentally defined domains likely reflect the inherent properties of ChIP-seq–based mapping, in which CenH3 enrichment frequently extends into surrounding pericentromeric chromatin [13, 29, 30].

In *N. crassa*, which possesses exceptionally large regional centromeres (∼150–400 kb), CentroFinder again localized compact centromeric regions (∼40–95 kb) positioned entirely within the experimentally defined centromeric spans for all chromosomes with available reference data (Table 1). Because *N. crassa* centromeres span exceptionally large genomic intervals, the fully nested CentroFinder predictions yield large apparent offsets (∼253–341 kb; median ∼277 kb) when measured relative to EDC boundaries. These offsets therefore reflect centromere size differences rather than prediction mislocalization, and instead indicate accurate identification of compact centromeric cores within expansive regional centromeres.

Together, these benchmarking results demonstrate that CentroFinder accurately localizes compact centromeric cores across diverse fungal centromere architectures, even when experimentally defined centromeric domains span hundreds of kilobases.

### 3.3 Integrated Multi-Feature Scoring Robustly Delineates Centromeric Cores

Across all benchmarked fungal genomes, individual centromere-associated features— including GC depletion, enrichment of transposable elements and tandem repeats, reduced gene density and CpG methylation, and localized long-read coverage anomalies—showed heterogeneous distributions along chromosomes. When considered independently, each feature frequently marked multiple genomic regions, particularly in repeat-rich or structurally complex backgrounds, and was therefore insufficient on its own to uniquely identify centromeric loci. In contrast, integrating these features into a composite centromere score resulted in the convergence of multiple centromere-associated signals at a single internal locus per chromosome. This convergence produced a pronounced and isolated maximum in the integrated CentroFinder score,enabling unambiguous identification of one centromeric region per chromosome (Figure 1). Notably, this pattern was consistently observed across species with markedly different centromere sizes, repeat compositions, and chromatin architectures (Table S1).

The effectiveness of this integrative strategy was particularly evident in genomes with large or degenerate centromeres, such as *N. crassa*, which has extensive repeat-rich regions spanning hundreds of kilobases. In these cases, individual features alone broadly marked pericentromeric domains, whereas their combined signal sharply localized compact centromeric cores within the larger experimentally defined regions (Table 1). Similarly, in *M. oryzae*, integration suppressed false positives in subtelomeric and repeat-dense regions, yielding a single dominant centromeric interval per chromosome despite widespread repeat content.

Together, these results demonstrate that robust centromere prediction in fungi requires a multi-feature framework that captures the combinatorial nature of centromeric identity. By jointly evaluating complementary sequence- and long-read–derived signals rather than relying on any single defining feature, CentroFinder achieves high specificity and consistency across diverse fungal genomes, accurately delineating conserved centromeric cores even within expansive or structurally complex centromeric domains.

## 4 Discussion

Accurate identification of centromeres remains a fundamental challenge in genome analysis due to their rapid sequence evolution, high repeat content, and lack of conserved defining motifs [6, 9, 29]. This challenge is particularly pronounced in fungi, where centromere architectures span an exceptional spectrum of sizes, sequence compositions, and epigenetic configurations even among closely related species [16, 17, 31, 32]. In this study, we introduce CentroFinder, a fungal-specific computational framework that integrates multiple genomic and long-read–derived features to enable robust, *de novo* centromere prediction from genome assemblies.

A central insight underlying CentroFinder is that no single genomic signal is sufficient to reliably define fungal centromeres across lineages. Features such as repeat enrichment, GC depletion, gene absence, DNA hypomethylation, or read-depth irregularities can each occur elsewhere in the genome, particularly in subtelomeric or structurally dynamic regions. By explicitly integrating these heterogeneous signals within a biologically informed weighted scoring framework, CentroFinder identifies the unique internal locus on each chromosome where centromere-associated features converge most strongly. This integrative strategy enables the robust discrimination of centromeric regions from other repetitive or compositionally biased loci.

Importantly, CentroFinder generalizes across fungi with markedly different centromere architectures. Despite substantial differences in centromere size, repeat composition, and chromatin organization among *C. deuterogattii, M. oryzae*, and *N. crassa* [27-29], CentroFinder consistently localized a single centromeric region per chromosome that overlapped experimentally defined centromeres. This robustness highlights the advantage of a feature-integration strategy that is agnostic to specific repeat families or species-restricted motifs—an essential property given the rapid evolutionary turnover of centromeric DNA in fungi.

Notably, CentroFinder predicts compact centromeric intervals that are fully contained within broader experimentally defined centromeric domains. This outcome reflects fundamental differences between sequence-based prediction and chromatin-based centromere mapping rather than systematic underprediction. Chromatin immunoprecipitation assays such as CENP-A or CenH3 ChIP-seq typically identify extended enrichment domains that include both centromeric and pericentromeric chromatin, particularly in species with large regional centromeres [13, 29, 33, 34]. In contrast, CentroFinder preferentially delineates the centromeric core—defined here as the interval where multiple centromere-associated sequence and long-read features co-localize most strongly—providing a practical computational proxy for functional centromere localization when experimental data are unavailable. This distinction explains the larger apparent positional offsets observed in species such as *N. crassa*, where expansive centromeric domains encompass a relatively compact functional core.

CentroFinder does not strictly require fully complete chromosome-level assemblies to accurately delineate centromeric cores. This is exemplified by *M. oryzae*, for which experimentally defined centromeres were originally mapped using a partially assembled genome, yet CentroFinder precisely localized centromeric regions overlapping these CENP-A–defined domains. As long as long-read sequencing data provide sufficient contiguity across internal chromosomal regions, the integration of multiple centromere-associated features remains effective. The principal limitation in more fragmented assemblies arises near chromosome ends, where incomplete representation of subtelomeric regions may complicate the exclusion of telomere-proximal signals rather than impair centromere detection itself.

In summary, CentroFinder provides a robust, interpretable, and biologically grounded framework for centromere prediction tailored specifically to fungal genomes. By integrating complementary genomic and long-read–derived signals, it overcomes the limitations of single-feature approaches and enables reliable centromere annotation across diverse fungal lineages. Because CentroFinder relies exclusively on sequence- and read-based information, its accuracy scales with assembly continuity and long-read coverage, making it well suited to modern long-read sequencing projects. We anticipate that CentroFinder will facilitate comparative studies of centromere evolution, karyotype dynamics, and genome structural variation, and will serve as a foundational tool for centromere-aware fungal genomics in the long-read era.

## Acknowledgments

The authors acknowledge the Research Computing and Data division at Tennessee Tech University (RRID:SCR_027555) for providing computational resources and support services that have contributed to the research results reported within this paper. Specifically, computations were supported by the National Science Foundation under Award #2127188. The authors also thank Robert H. Proctor (U.S. Department of Agriculture) for helpful comments and Michael Freitag for providing *Neurospora crassa* experimental data.

---

**Table S1.Chromosome-level comparison between CentroFinder-predicted centromeres and experimentally defined centromeres in benchmark fungal genomes.** For each chromosome, genomic coordinates and lengths of centromeres predicted by CentroFinder are reported alongside experimentally defined centromere (EDC) coordinates derived from CENP-A or CenH3 mapping. The offset column indicates the absolute positional difference (in base pairs) between the boundaries of the predicted centromeric interval and the corresponding EDC. Results are shown for **(A)** *Cryptococcus deuterogattii* R265, **(B)** *Magnaporthe oryzae* Guy11, and **(C)** *Neurospora crassa* OR74A.

## Notes

### Competing Interest Statement

The authors have declared no competing interest.

